# Assessing the relationship between female aggression and male phenotype in two sister *Malurus* fairywren species

**DOI:** 10.1101/2023.10.19.563140

**Authors:** John Anthony Jones, William E Feeney, Darryl N Jones, Doka Nason, Serena Ketaloya, Jordan Karubian

## Abstract

In a large and ever-growing number of animal species, it is now appreciated that females use colors as a visual signal in a range of social interactions, including both courtship and territorial aggression. Yet, it remains unclear whether female color phenotypes and/or aggressive behaviors are correlated with any attributes of their mate’s phenotype. For example, we might expect species in which males contribute more to parental care or territorial defense to have more colorful or aggressive females. On the other hand, within species, we might expect those females mated to higher quality males to be more colorful or aggressive than those mated to lower quality males. To begin to address these possibilities, we conducted a preliminary study in two sister taxa of fairywren (Maluridae) with distinct life-history strategies and plumage dichromatism: white-shouldered fairywrens (*Malurus alboscapulatus moretoni*) in tropical Papua New Guinea, a species in which both males and females have ornamented plumage and jointly defend territories year-round, and red-backed fairywrens (*M. melanocephalus melanocephalus*) in temperate Australia, a sexually dichromatic species with ornamented males and unornamented females. At the between species level, we predicted white-shouldered fairywren females would be more aggressive in same-sex interactions than red-backed fairywrens, as both white-shouldered males contribute to year-round territorial defense, whereas territories break-down during non-breeding in red-backed fairywrens. Further, we predicted that, within species, females mated to males of higher quality would be more aggressive in simulated same-sex encounters. Between species, female white-shouldered fairywrens were more aggressive on average than female red-backed fairywrens as predicted. Within both species, indices of male quality were not related to female aggression (although there was a non-significant tendency for more aggressive female white-shouldered fairywren to have heavier mates with longer tails). These results point to a need for additional research exploring relationships between life history, female plumage, and female aggressive behaviors in a wider range of species.

## Introduction

Social selection theory posits that competition over any critical resource, whether sexual or non-sexual in nature, should lead to the evolution of competitive traits (e.g., aggression, ornaments, armaments, etc.) that aid in resource acquisition (Crook 1972; West-Eberhard 1979; Lyon and Montgomerie 2012; Tobias et al. 2012). Though the study of competitive interactions has been well explored both theoretically and empirically from the male perspective, female animals also compete aggressively for access to limited resources crucial to survival and reproduction (Langston et al. 1990; Slagsvold and Lifjeld 1996; Fedy and Stutchbury 2005; Rosvall 2008). Despite centuries of academic neglect, there is now little doubt that female-female competitive interactions are adaptive and undirect selection pressures in a variety of taxa (Clutton-Brock 2007; Stockley and Bro-Jørgensen 2011; Tobias et al. 2012; Cain and Ketterson 2013; Stockley and Campbell 2013; Cain and Rosvall 2014; Lipshutz and Rosvall 2021; Fischer et al. 2023). Indeed, social selection is often invoked as an explanation for the evolution of competitive phenotypes in females, as the most aggressive females are those who are more likely to acquire resources necessary for breeding (Yasukawa and Searcy 1982; Jawor et al. 2006; Pryke 2007; Rosvall 2008), ultimately achieving greater reproductive success (Slagsvold and Lifjeld 1996; Rosvall 2011; Cain and Ketterson 2012).

Because selection maximizes reproductive success in each sex differently (Trivers 1972; Westneat and Sargent 1996; Kappeler et al. 2022), it remains unclear if the same ecological and social stimuli that commonly predict patterns of male aggression equally apply to females. Would we expect to find the most aggressive females to be mated to the highest quality males? Females are typically not limited by the number of potential mates available, and as such, a common perception is such that females-female competition most often occurs over ecological resources that enhance and/or are necessary for reproductive success (e.g., food availability, oviposition sites, etc.) rather than mates per se (Heinsohn et al. 2005; Stockley and Bro-Jørgensen 2011; Tobias et al. 2012; Stockley and Campbell 2013). Nevertheless, while males often compete for access to the greatest number of potential mates, females may compete for access to the highest quality mates that provide the best parental care, indirect (genetic) benefits, and/or access to highly preferred nesting locations (Rosvall 2011). For example, intrasexual aggression among female sharknose gobies (*Elacatinus evelnae*) is most intense over larger males, as this phenotype is typically associated with elevated paternal care (Whiteman and Côté 2003). However, intrasexual female competition over the highest quality mates is not universal; aggression among female dark-eyed juncos (*Junco hyemalis*) is most intense over nonsexual resources (nest site quality), with male quality as a secondary consideration (Cain 2014). Given this overall paucity and diversity of findings on degree of within species female competition, such as whether female aggression or other traits accurately predict quality of their mates, further explorations would help to advance our understanding of female signal evolution.

On a similar note, a comparison of closely related species that differ in the relative degree of female social competition may help elucidate the role that such variation in the competitive environment and/or overall life-history plays in shaping the intensity of female aggression. Variation in the breading season length, mating system, and food resources all have a large role in influence the direction of territorial aggression among songbirds (Stutchbury and Morton 2023) including females. The strength of social selection acting on males and females is predicted to be similar in tropical regions where sex role convergence in year-round territory defense and parental care is common (Kunkel 1974; Slater and Mann 2004; Stutchbury and Morton 2023) versus that of temperate regions, where territory defense typically occurs only in the spring and summer (Fedy and Stutchbury 2005; Catchpole and Slater 2008). Tropical species often consist of both sexes expressing some form of mutual ornamentation (Karubian 2013; Dale et al. 2015), potentially reflecting how shared phenotypes function as signals in mate-choice and/or competitive contexts in both sexes (Kraaijeveld et al. 2007; Murphy et al. 2009). In contrast, species occupying temperate ecosystems are commonly dichromatic, with males stereotypically highly adorned and territorial, whereas females are often muted in plumage coloration and do not defend the territory as vigorously (Stutchbury and Morton 2023). These observations are consistent with the idea that female-female competition experienced in tropical regions may exceed that commonly experienced by their temperate congeners, but there are limited examples of comparative behavioral studies between closely related, recently diverged species. Moreover, few behavioral assays reliably compare aggression between geographically isolated species while controlling for the influence of the social environment.

In this descriptive study, we studied two sister species that vary distinctly in overall life history strategy and sexual dichromatism: (1) white-shouldered fairywrens (*Malurus alboscapulatus moretoni*), a tropical songbird endemic to New Guinea that defend territories year-round, are mutually ornamented (in all but one subspecies (Enbody et al. 2019)), and have convergent sex roles, and (2) red-backed fairywrens (*M. melanocephalus melanocephalus*) of subtropical-to-temperate Australia, a dichromatic species with seasonal territoriality and breeding (Fig. 1). Both species are socially monogamous, but differ markedly in overall sociality and reproductive strategy. Red-backed fairywrens are highly social, with nesting groups that consist of breeding pair and often (though not always) contain additional auxiliary (both kin and non-kin) helpers (Rowley and Russell 1997; Karubian 2002; Webster et al. 2008), whereas helpers are the exception, rather than the norm, in the subspecies of white-shouldered fairywrens of the current study (Enbody et al. 2019). White-shouldered fairywrens also have lower levels of extra-pair paternity (∼33%) and have smaller scaled cloacal protuberance volumes on average (7.85 mL^3^) than red-backed fairywrens (∼56% and 16.94 mL^3^, respectively (Brouwer et al. 2017; Enbody et al. 2019)); among *Malurus*, larger cloacal protuberance volumes is positively associated with testes size, sperm quality and overall rates of extra-pair paternity (Tuttle et al. 1996; Rowe and Pruett-Jones 2011; Rowe and Pruett-Jones 2013). Finally, male red-backed, but not white-shouldered fairywrens, exhibit delayed plumage maturation, such that the majority of males are unornamented and appear identical to females (to humans, but not birds; Karubian et al. 2008) during their first breeding season (Webster et al. 2008; Karubian et al. 2011). There is evidence that male coloration is sexually selected in red-backed fairywrens, as experimental research between red-backed fairywren subspecies have shown that females prefer redder dorsal plumage over orange coloration, with redder males siring significantly more extra-pair young (Baldassarre and Webster 2013).

**Figure 1.**
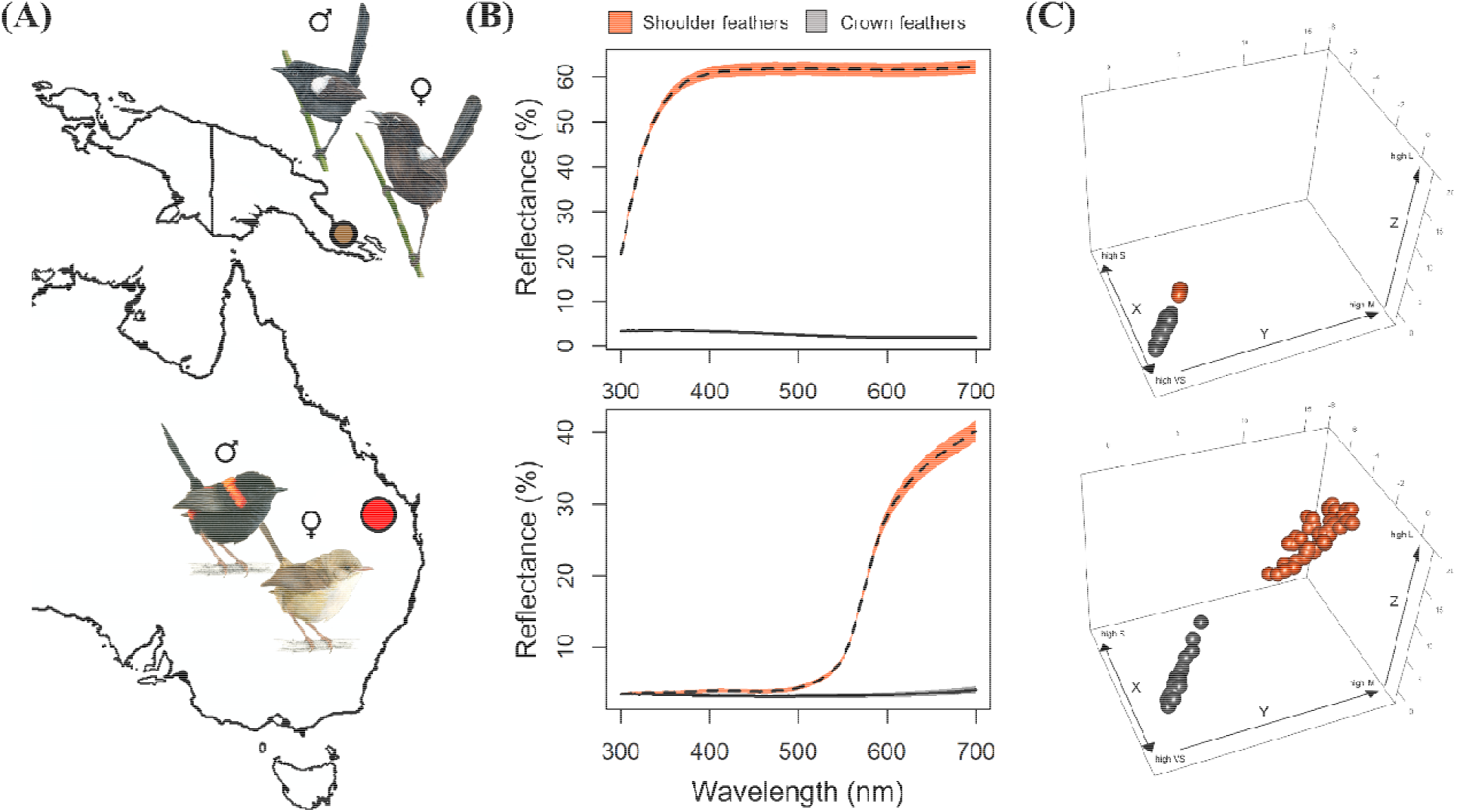
Visual illustration of (A) approximate sampling location and (B, C) male plumage coloration of white-shouldered fairywrens of New Guinea (top) and red-backed fairywrens of Australia (bottom). (B) Plumage reflectance spectra showing mean (± SD) amount of light reflected at a given wavelength of shoulder (orange, dashed lined) and crown (gray, solid line) feathers. (C) Variation in plumage coloration represented in avian tetrahedral colorspace *XYZ* coordinates, set to the same scale for species comparison purposes. Fairywren illustrations provided by Allison Johnson.

Based on these species differences in mating systems and life-history strategy, we predicted that female white-shouldered fairywrens would be more likely to experience intense selection pressure via intrasexual competition than female red-backed fairywrens, and thus display overall higher levels of aggression. Both within and between species, we quantified variation in how individuals respond to same-sex intruders through the use of mirror image simulations behavioral assays that put individuals up against their own mirrored reflection, mimicking an intrasexual challenge against a rival of equivalent visual quality (Gallup Jr 1968; Leitão et al. 2019; Jones, Boersma, Liu, et al. 2022). A strength of this behavioral approach is that it allows for reliable and relatively standardized comparisons between two species of fairywrens that differ in mating strategy, life-history, and even geographical location. Moreover, these assays isolate focal individuals from their social mate, a consideration of particular noteworthiness for species which engage in joint territory defense like *Malurus* (Jones, Boersma, and Karubian 2022). We additionally conducted an exploratory analysis on if more (or less) aggressive females are partnered with males of higher phenotypic quality by correlating aggression scores with male plumage coloration and body condition indices. We predicted that, if females compete over access to the most attractive males, the most aggressive females will be partnered with males of higher phenotypic quality. This preliminary assessment of how female aggression may relate to between species variation in life history and dichromatism, and within species variation in mate quality, is intended to promote and guide further research rather than a definitive test of these predictions per se.

## Methods

### Study species and general field methods

Both sister species are predominately grassland specialists, but differ significantly in mating system structure, overall life-history strategy, and if females are ornamented or not (Rowley and Russell 1997). In New Guinea, white-shouldered fairywren plumage coloration remains consistent year-round in both sexes (Fig. 1). Male plumage coloration is consistently black-and-white throughout the island, but females vary allopatrically in plumage phenotype that tracks with subspecies (Rowley and Russell 1997; Enbody et al. 2019). The current study is of *Malurus alboscapulatus moretoni*, a subspecies where both sexes are mutually ornamented, with such female coloration having been derived from that of an unornamented phenotype that resembles their sister species (Enbody et al. 2019; Enbody et al. 2022). In contrast, male red-backed fairywrens in Australia seasonally molt between an ornamented plumage phenotype (i.e., eumelanin-based black body coloration and carotenoid-based red backs and shoulders) during the breeding season and a drab (phaeomelanin-based brown) phenotype that resembles that of females during the non-breeding season (Rowley and Russell 1997; Karubian 2002; Karubian 2008). Female plumage color typically does not vary, such that they remain unornamented year-round, although it is possible that variation in bill coloration may exist as it does in males (Karubian 2008).

We studied a population of white-shouldered fairywrens in Podagha Village, Milne Bay Province, Papua New Guinea (9.692□S, 149.895□E) from June-July 2019, and red-backed fairywrens in Samsonvale, Queensland, Australia (27.271□S, 152.857□E) from August-November 2019. We captured individuals with mist-nets for banding and feather collection. To explore for evidence that female coloration is related to male phenotype, we focused on three standard body measurements that we suspected may be important differences between sexes and/or species: (1) mass (± 0.01 g): both species exhibit size dimorphism, such that males are slightly heavier than females (Rowley and Russell 1997); (2) Tail length (±0.01 mm): shorter tails are thought to be a signal of social dominance in *Malurus* (Swaddle et al. 2000; Karubian et al. 2009); (3) Scaled cloacal protuberance (CP) volume (0.01 mL^3^; Tuttle et al. 1996; Rowe and Pruett-Jones 2011, 2013). We calculated scaled CP as the ratio of CP volume (calculated as: 3.14 * (CP height/2) * (CP width/2) * CP length) to body mass for males (Tuttle et al. 1996).

### Mirror image stimulation

We performed mirror image stimulation assays in both sexes of both fairywren species following the methods of Jones et al. (2022a,b). As most passerine species appear to be unable to self-recognize (Kraft et al. 2017, but see Prior et al. 2008), we interpreted aggressive behaviors as a response to a perceived same-sex rival of equivalent visual quality. We conducted our mirror assays from 0600-1100 or 1530-1730 local time (GMT +10) while avoiding rain and the intense midday heat. We held birds for ≥15 min post-capture, but prior to mirror assays, as this is when circulating corticosterone has likely reached its asymptotic peak and is unlikely to influence behavior beyond the scope of the current study (Cockrem and Silverin 2002). We temporarily placed fairywrens in a cage measuring 60 cm (length) x 40 cm (width) x 40 cm (height). One side of the cage consisted of a mirror that was initially hidden by a wooden cover, as well as three perches at the same height, but at uniform distance classes away from the mirror (defined as “close”, “neutral”, or “far” from the mirror; see Fig. 1C *in* Jones et al. 2022a). We covered the experimental cage on each side with a white cloth except the one exposed to the video recorder to (1) limit external stimuli influencing individual behavior and (2) reduce the number of potential exits that might distract the bird from the mirror. We placed the cage on the ground within the focal bird’s territory in a shaded location. We allowed focal individuals 5 min to acclimate to the cage prior to exposure to their mirrored reflection for ∼7 min. In the current study, we report the behavioral responses post-mirror exposure, as this is the only period in which focal birds displayed aggressive behaviors (see also Jones et al. 2022a). We scored the relative proportion of time spent at each distance class relative to the mirror (e.g., close to the mirror is indicative as a more aggressive stance than far from the mirror) and calculated the total instances of aggression observed (i.e., physically striking the mirror, threatening displays, and soft songs). We video recorded the mirror assays using a partially camouflaged Sony HDR-CX405 Handycam (Tokyo, Japan).

### Plumage coloration analysis

In white-shouldered fairywrens, our *a priori* expectation was that achromatic coloration (i.e., brightness, or percent of light reflected) and not chromatic variation, of the crown (melanin-based black feather coloration) and the shoulder (structurally-based white) feather regions were most likely to serve a signaling function. This assumption is based, in part, to the fact that crown feather coloration is the most variable black feather in this species (Jones unpubl. data). Nonetheless, because black plumage regions in male plumage are iridescent (whereas females are a matte black), we did explore the chromatic variation of these regions as well, although there is exceptionally little chromatic variation among males (Fig. 1). For male red-backed fairywrens, our *a priori* expectation was that the chromatic variation of the red/orange shoulder (carotenoid-based, Rowe & McGraw 2008) would be important to consider, as these carotenoid colors are targets of sexual selection (Karubian 2002; Karubian et al. 2008; Webster et al. 2008; Baldassarre and Webster 2013). For consistency, we also explored the black crown coloration in red-backed fairywrens.

We collected ≥6 feathers from each feather region from each male captured and stored them in a cool, dark environment until spectral analysis (detailed methods available in Jones et al. 2022a). We taped feathers on black cardstock (base reflectance: ∼10%) in a manner that resembled the way feathers naturally lie on a bird. We then made three repeated measures of reflectance for each feather patch using a USB2000+ spectrometer (R400-7-UV-VIS probe, RPH-1 probe holder) with a PX-2 pulsed xenon light source using OceanView software (v.1.6.7; Ocean Optics, Dundin, FL, USA). The resulting reflectance spectra generated were calibrated relative to a white standard that reflects 100% of light evenly from 300-700nm (Ocean Optics WS-2). We took the mean of the three spectra readings and binned them into 5 nm increments to be used in a psychophysical model of avian vision (Vorobyev et al. 1998; Vorobyev and Osorio 1998), which explores color variation from the bird’s visual perspective (as implemented by Delhey et al. (2015)).

Birds possess four types of single cones in their retinas that are sensitive to and detect variation in chromatic coloration (very short (VS), short (S), medium (M), and long (L) wavelengths) and one type of double cone responsible for achromatic (i.e., brightness) variation sensitivity (Cuthill 2006). White-shouldered and red-backed fairywrens are both V-type species (Ödeen et al. 2012) – species that possess ability to perceive UV wavelengths but with lower sensitivity than U-type species (Delhey et al. 2013; Ödeen & Håstad 2013)). Thus, we used the mean V-type peak sensitivity 416, 478, 542, 607 nm (Endler and Mielke 2005). We calculated signal-to-noise ratios for each cone type using formula 10 in Vorobyev et al. (1998), using the mean proportion of cones present in V-type birds from Hart (2001: VS = 0.381, S = 0.688, M = 1.136, L = 1.00), a Weber fraction of 0.1 for the L cone (Olsson et al. 2018), and an irradiance spectrum of standard daylight (D65; Vorobyev et al. 1998). This model of avian vision reduces variation among reflectance spectra to a set of three chromatic coordinates that define their position in avian tetrahedral colorspace: “*X*”, “*Y*”, and “*Z*.” Here, the *X* axis represents the relative stimulation of the S cone relative to the VS cone, *Y* represents the relative stimulation of the M cone to both the VS and S cones, and *Z* represents the relative stimulation of the L cone to the VS, S, and M cones. Distances between points in tetrahedral colorspace are reported as Just Noticeable Differences (JNDs); values representing >1 are considered distinguishable.

We summarized *XYZ* coordinates with a separate principal components analysis (PCA) per plumage region per species using a covariance matrix (*sensu* Jones et al. 2022). All PCAs resulted in one component (hereafter: PC1_chroma_) that explained >92% of the variation (Table 1). For melanin-based black crown coloration, both species had intermediate values for the *X* axis and larger, negative values for both *Y* and *Z*; spectra with larger, more negative Crown-PC1_chroma_ values for this feather region provided low stimulation of the L cone relative to VS + S + M cones, low stimulation of the M cone relative to VS + S cones, and higher stimulation of the VS cone relative to the S cone. Positive Crown-PC1_chroma_ values were those richer in shorter wavelengths (UV/blue) than in longer wavelengths (red). In white-shouldered fairywrens, higher Shoulder-PC1_chroma_ values stimulated the same cones as crown coloration (as both colors are achromatic), although with a greater stimulation of VS than S cones. However, the eigenvalues for both crown and shoulder coloration in white-shouldered fairywrens were <1.0 despite comprising a vast majority of the variation (>92%), suggesting that there is minimal chromatic variation in these plumage regions. For the carotenoid-based shoulder colors found in red-backed fairywrens, positive Shoulder-PC1_chroma_ scores were feathers that provided higher stimulation of the L cone relative to VS + S + M cones and higher stimulation of the M cone relative to VS + S. Thus, individuals with larger values along this PC axis were individuals with richer red coloration and this axis explained little variation in the UV/blue region. We calculated overall brightness of both the crown and scapular feather patches by calculating the achromatic contrast between each plumage patch and an exceptionally dark spectrum by setting the double cone quantum catch value to 0.001 (Delhey et al. 2015).

**Table 1.** Loadings for principal components analysis of *XYZ* coloration (PC1_chroma_) for male red-backed and white-shouldered fairywrens. Loadings are each feather patch per species for a single component which explains the majority of variation across patches.

|  | White-shouldered fairywren |  | Red-backed fairywren |  |
| --- | --- | --- | --- | --- |
|  | Crown | Shoulder | Crown | Shoulder |
| Eigenvalue | 0.470 | 0.014 | 2.950 | 13.51 |
| Variation explained (%) | 0.928 | 0.972 | 0.984 | 0.956 |
| *X* | 0.367 | 0.677 | 0.389 | -0.005 |
| *Y* | -0.638 | -0.607 | -0.633 | 0.727 |
| *Z* | -0.677 | -0.416 | -0.669 | 0.687 |

### Statistical analysis

We performed all statistical analyses in R v.4.1.1 (R Core Team 2021). Following a previously established method for mirror image stimulations (Leitão et al. 2019; Jones, Boersma, Liu, et al. 2022), we aggregated mirror strikes, pecks, vocalizations, and displays in response to the focal individual’s mirrored reflection by summing them into one overall “Aggression” value, as the latter behaviors were infrequent on their own (Jones, Boersma, Liu, et al. 2022). Using these data alongside the proportion of time spent near versus far from the mirror, we explored species differences in female aggression via a Welch’s t-test (as variance was unequal between species). Next, we generated a composite index of aggression in response to their mirrored image with separate PCAs for each species and sex individually (Leitão et al. 2019; Jones, Boersma, Liu, et al. 2022). Behavioral responses during mirror image stimulation assays resulted in a single component with an eigenvalue >1.0 to be retained for further analysis (sensu Leitão et al. 2019; Jones et al. 2022b; Table 2). All behavioral PCs loaded similarly, regardless of sex or species, such that individuals with higher PC1_MIS_ loading scores were birds who responded more aggressively towards their mirror reflection, indicative of a greater proportion of time near the mirror rather than far away from it coupled with increased attack rate.

**Table 2.**
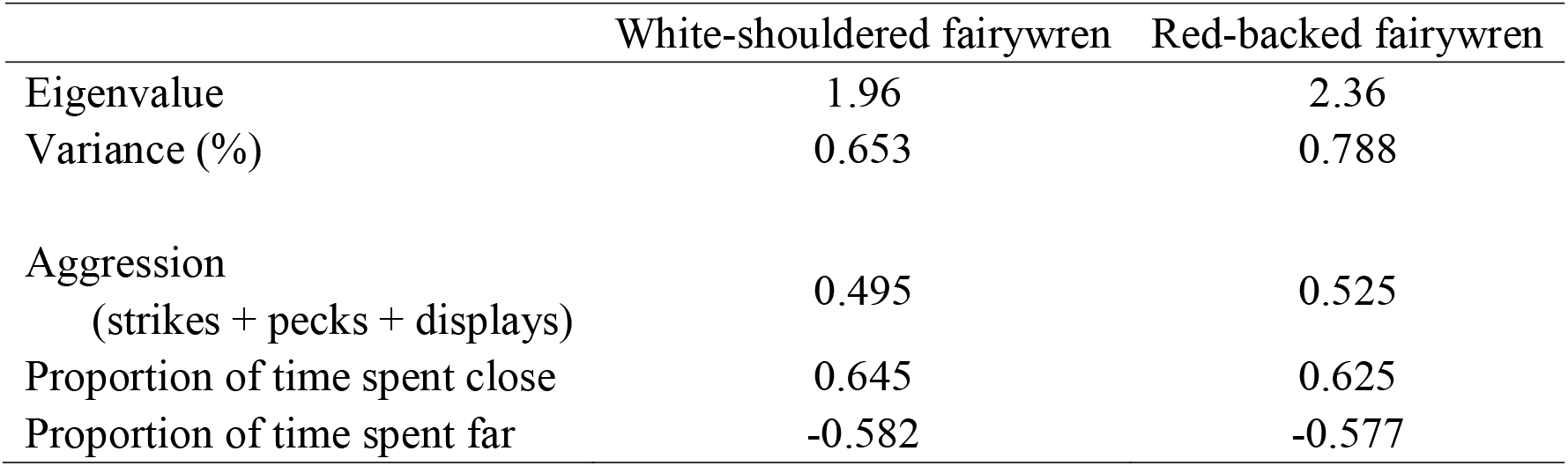
Loading scores for the principal components analysis (PC1_MIS_) exploring how fairywrens respond to mirror image stimulation.

To explore the relationship between male phenotypic quality and female aggression, we ran a series of multiple regression analyses coupled with a finite sample corrected Akaike’s information criterion (AICc) model selection procedure. For each model (i.e., species), individual female aggression scores were set as a dependent variable and our fixed effects were seven (putative) indices of male condition (center and scaled tail length, mass, scaled CP volume, as well as chromatic and achromatic chroma of both the crown and scapular feather patches); only final models are presented in the main text (but see Supplemental Material for full AICc table). Finally, we ran a separate, complementary statistical analysis using the same male morphological traits, condensed via PCA (following the approach of Cain (2014)). The results between statistical approaches were consistent and we present these findings in the Supplemental Material.

### Ethical note

Our study was carried out in strict accordance with the guidelines established by the Tulane University Institutional Animal Care and Use Committee (#0395R2), Griffith University’s Animal Ethics Committee (ENV/08/19/AEC), and in adherence to research permits from the Conservation and Environment Protection Authority of Papua New Guinea (#99902100765). All birds were captured, processed, exposed to mirror assays, and released in under one hour. We continuously monitored our mist-nets and removed birds immediately upon hitting the net. All individuals involved in removal of birds from mist-nets were trained in the appropriate technique to extract and safely handle birds.

## Results

### Species comparison

We conducted mirror assays on 26 female red-backed fairywrens and 44 white-shouldered fairywrens. On average, white-shouldered fairywren females were more aggressive than red-backed fairywren females, determined as a greater rate of aggressive behaviors observed per min (t = 3.53, df = 64.40, p < 0.001; Fig. 2) and proportion of time near the mirror (t = 3.61, df = 45.32, p <0.001). Red-backed fairywrens were more likely to be positioned far from the mirror than white-shouldered fairywrens (t = 3.68, df = 34.05, p < 0.001).

**Figure 2.**
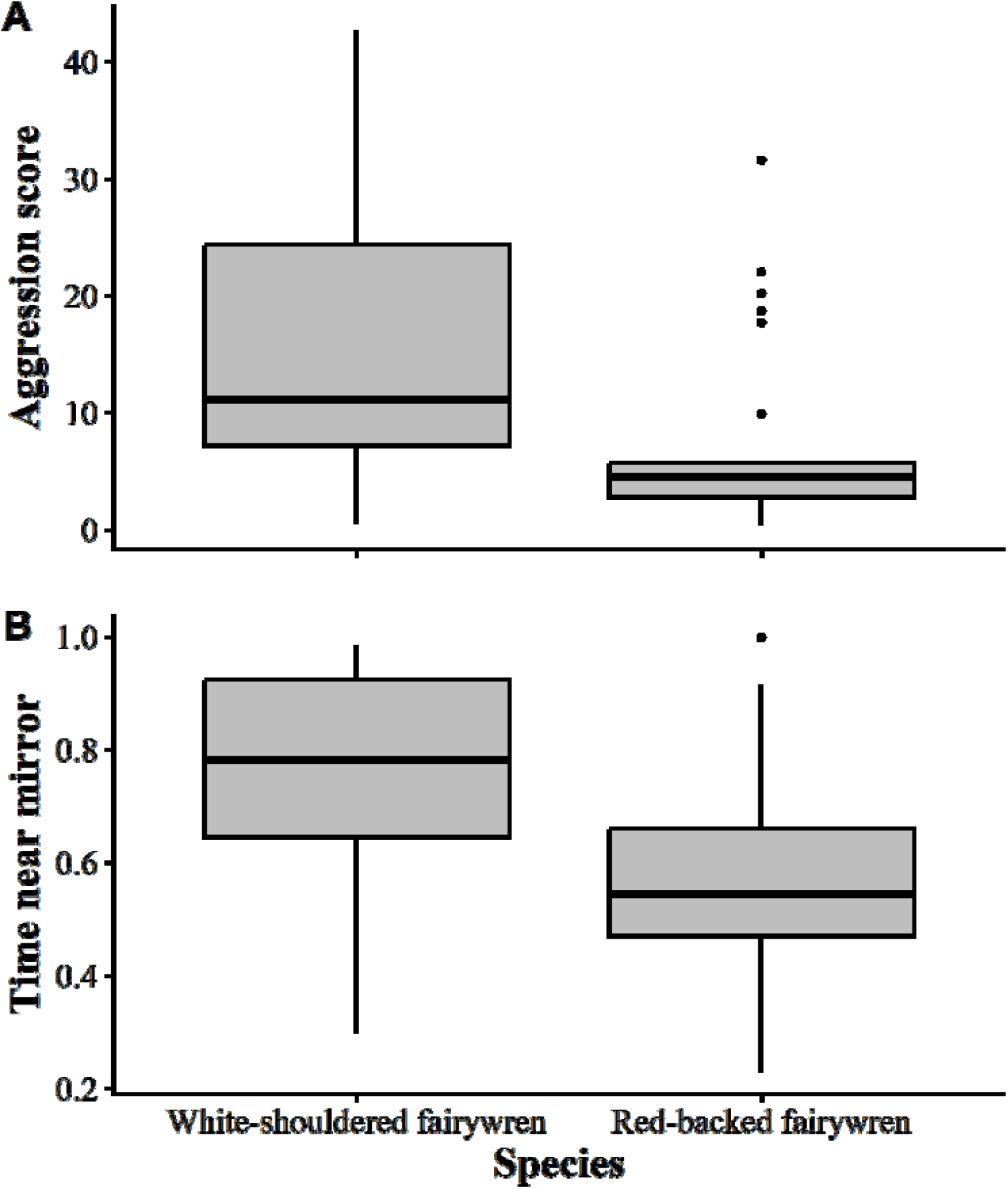
Responses of white-shouldered fairywrens and red-backed fairywrens to mirror image stimulation. (A) Aggression score represents the aggregation of mirror strikes, pecks, songs, and displays preformed throughout the mirror assay. On average, female white-shouldered fairywrens were significantly more aggressive towards their reflection than red-backed fairywrens, frequently attacking the mirror. (B) Time near mirror represents a proportion of time relative to the length of the trial, such that white-shouldered fairywrens spent a greater proportion of time near the mirror compared to red-backed fairywrens.

### Female aggression with respect to male phenotypic quality

Previous work in this population of white-shouldered fairywrens revealed that female aggression is unrelated to her own plumage coloration and body size (Jones, Boersma, Liu, et al. 2022). We tested the relationship between male mate quality attributes on the aggression of 38 individual female white-shouldered fairywrens. The best fitting model predicting female white-shouldered fairywren aggression included male mass, tail length, and CP volume, although the overall model was not statistically significant (multiple r^2^ = 0.10, F_3, 37_ =2.45, p = 0.08; Fig. 3; Table 3; Table S1 for full model selection parameters). Within the best fitting model, females were most aggressive if their mate tended (non-significantly) to be heavier (β est. = 0.62, p = 0.10) and have a longer tail (β est. = 0.42, p = 0.06); despite appearing in the top model, CP volume was not significantly related to aggression (β est. = 0.04, p = 0.50). There were an additional five models < 2 ΔAICc (Table 3), all of which contained some combination of the above three characteristics and all in the same direction

**Table 3.**
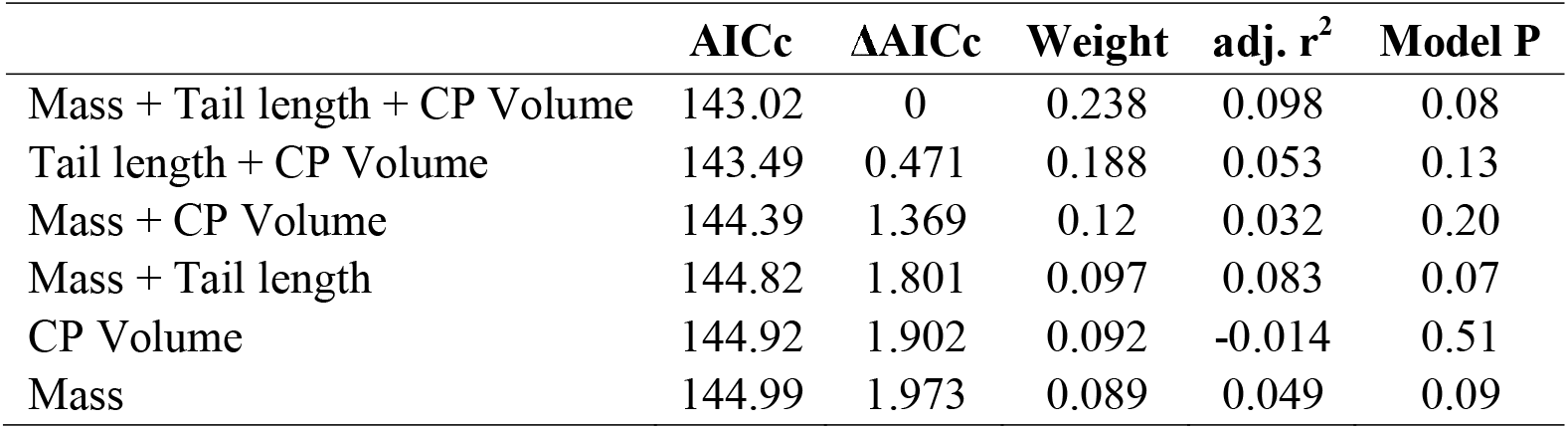
Abbreviated model selection table exploring which male phenotypic variables best predict aggression in female white-shouldered fairywrens. Presented are the models < 2 ΔAICc from the best fitting model; see Table S1 for full model parameters.

| | AICc | $\Delta AICc$ | Weight | adj. $r^2$ | Model P |
| --- | --- | --- | --- | --- | --- |
| Mass + Tail length + CP Volume | 143.02 | 0 | 0.238 | 0.098 | 0.08 |
| Tail length + CP Volume | 143.49 | 0.471 | 0.188 | 0.053 | 0.13 |
| Mass + CP Volume | 144.39 | 1.369 | 0.12 | 0.032 | 0.20 |
| Mass + Tail length | 144.82 | 1.801 | 0.097 | 0.083 | 0.07 |
| CP Volume | 144.92 | 1.902 | 0.092 | -0.014 | 0.51 |
| Mass | 144.99 | 1.973 | 0.089 | 0.049 | 0.09 |

**Figure 3.**
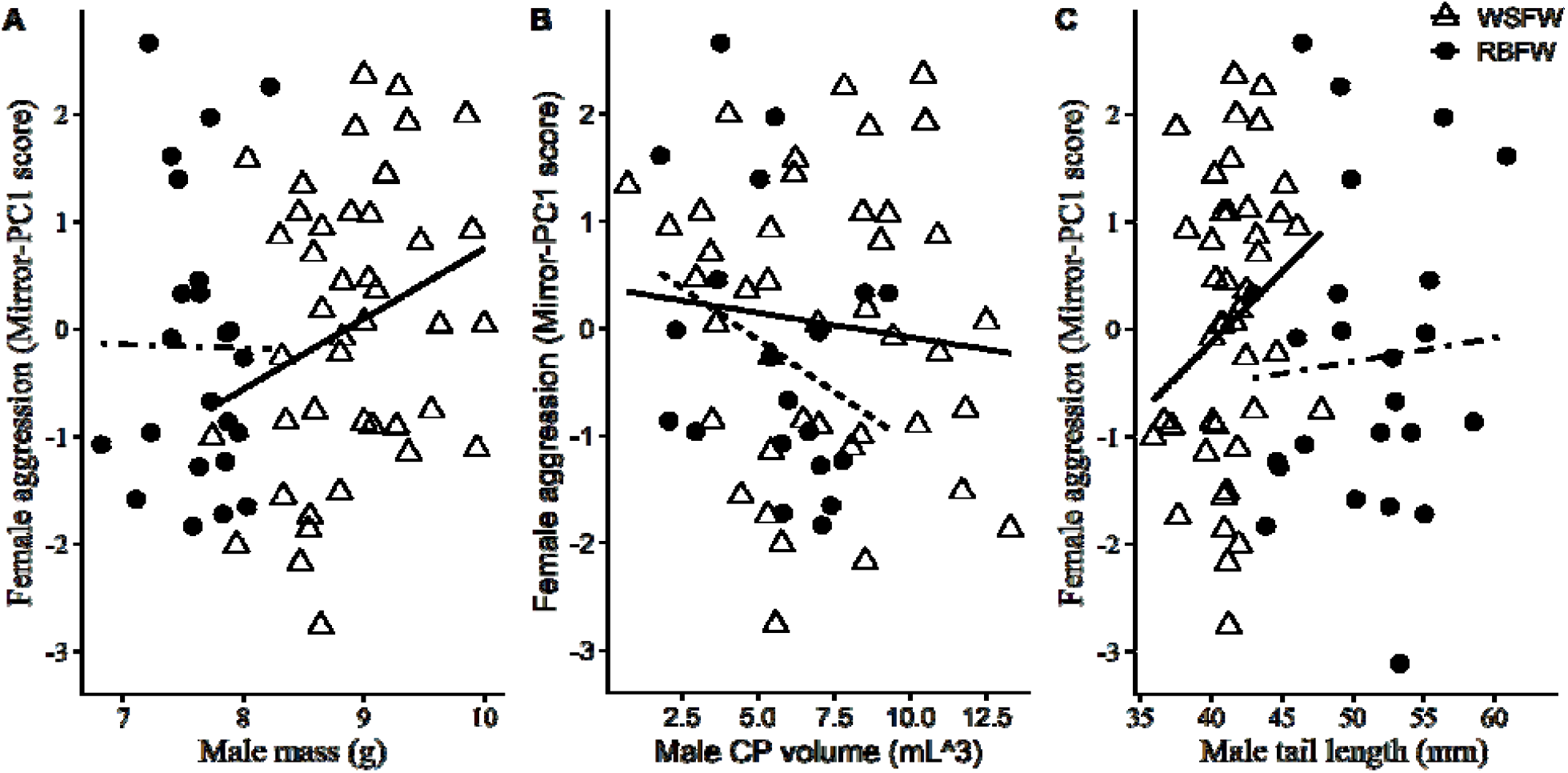
Relationship between female aggression and social mate (A) mass, (B) CP volume and (C) tail length in white-shouldered fairywrens (WSFW; triangle; solid line) and red-backed fairywrens (RBFW; circle, dotted line).

In red-backed fairywrens, we first ran a series of Pearson’s correlations and found that female aggression was not related to her own age (p = 0.34) nor any standard morphological measurement (e.g., mass, tarsus length, tail length; all p > 0.16). Including only pairwise observations, we compared female aggression to her mated male’s phenotype for 18 female red-backed fairywrens. The best fitting model contained center and scaled CP volume as the sole predictor, but this model was not statistically significant (multiple r^2^ = 0.09, F_1, 16_ =1.65, p = 0.22; Fig. 3; Table S2 for full model selection parameters). There was one additional model < 2 ΔAICc containing male mass alongside CP volume, but neither parameter were significant predictors of aggression (both p > 0.19).

## Discussion

We explored how females in two sister *Malurus* species with marked differences in mating system and overall life-history strategy vary in aggressive behaviors and whether or not females of these species who are mated with males in higher physical condition are more aggressive. First, we found support for our prediction that white-shouldered fairywrens, a tropical songbird species endemic to New Guinea that experiences intense year-round competition, would be more aggressive on average than their temperate sister species, red-backed fairywrens of Australia. Second, we failed to find a relationship between male coloration and female aggression in white-shouldered fairywrens. However, that male mass and tail length emerged in our analysis as potential (as they were marginally non-significant) predictors of female aggression is interesting and warrants further study; the females who were most responsive to mirror stimulation were mated to males who tended to be heavier, but also had longer tails. Nonetheless, as these results were not statistically significant, we failed to find evidence for a signaling function for these phenotypic traits and any other possibilities are at best speculative for now. In red-backed fairywrens we found that the best predictor of female aggression from our dataset was male CP volume, but critically, this relationship was not statistically significant nor was there an apparent trend. These results suggest that there is not a relationship between female red-backed fairywren aggression and any metric of male condition we measured, which may indicate that females do not compete for access to more attractive males in this species.

Though we cannot say with certainty what resource females compete over, our results suggest that, for both species, it is likely not for males of the highest phenotypic quality. A common assumption is that females preferentially mate with the most attractive male, though an appreciation for how females actively compete for such males has only occurred recently (reviewed in Rosvall 2011, Hare and Simmon 2019). For example, female topi (*Damaliscuss lunatus*) use cranial horns to attack rival females over access to preferred males (Bro-Jørgensen 2002). However, recent evidence suggests that intrinsic quality of one’s partner may not be as important for partnering decisions as previously thought, at least within socially monogamous systems (Ihle et al. 2015; Wang et al. 2017; Griffith 2019; Hurley et al. 2020). Rather, the quality of behavioral interactions between females and males (e.g., cooperativeness) in these species may be more important than phenotypic quality, particularly for tropical birds. Perhaps chick-rearing in the tropics requires the combined efforts of both parents, and as such, whether or not a male is physically attractive may only be useful to a female if male phenotype honestly signals parental investment (Préault et al. 2005; Pagani-Núñez and Senar 2014). In this sense, while male phenotypic quality may be important for initial pairing for some species (Griffith 2019), females in species that form longer-lasting pair ponds must also weigh the potential genetic quality of offspring (Hasselquist et al. 1996; Mays and Hill 2004), the fighting capacity (Reudink et al. 2009), and/or parental care (Préault et al. 2005; Pagani-Núñez and Senar 2014) provided when choosing a mate. This is consistent with our results, in that female intrasexual aggression was not related to any attribute of the mated male phenotype in either socially monogamous species. With that said, that male phenotype trended towards statistical significance in white-shouldered fairywrens may be noteworthy and warranting further study. It is possible that male phenotype is a secondary consideration for female was is in dark-eyed juncos (Cain 2014), though it also possible that the highest quality males disproportionately control the highest quality territory sites that females compete over.

In line with our expectation that the intensity of female-female aggression experienced in tropical regions would exceed that of temperate congeners, we found that female white-shouldered fairywrens are, on average, significantly more aggressive towards intrasexual rivals than red-backed fairywrens, their sister species with clear sexual dichromatism and female preferences for male coloration (Baldassarre and Webster 2013). In New Guinea, it may be that females perceive potential rivals as existential threats to their overall reproductive success, driving the behavioral patterns we observed. The strength of selection acting on males and females is predicted to be similar between sexes in tropical regions (Kunkel 1974; Slater and Mann 2004). Thus, the relative contribution one sex makes towards rearing offspring relative to their mate is likely to converge in socially monogamous tropical songbirds that pair and defend territories year-round (Stutchbury and Morton 2023). In this sense, how (dis)similar males and females are may act as a proxy for the strength or type of social selection acting on each sex independently. Several comparative studies have suggested that the female ornamentation is greatest closer to the equator (Karubian 2013; Dale et al. 2015), which likely reflect how shared phenotypes function as signals in mate-choice and/or competitive contexts in both sexes (Kraaijeveld et al. 2007; Murphy et al. 2009). In contrast, species that live in temperate ecosystems are often dichromatic, such that males are stereotypically highly adorned and territorial, whereas females are often muted in plumage coloration and do not defend the territory as vigorously (Stutchbury and Morton 2023). With that being said, Fargevieille et al. (2023) recently described how females of temperate songbird species with greater paternal care tend to be slightly brighter and more chromatic, suggesting that differences in dichromatism per se may not be a reliable indication of the strength of social selection.

An interesting distinction between our two focal species is the presence of auxiliary helpers. Most fairywren species are thought to breed cooperatively, with groups comprised of a breeding male and female alongside kin and non-kin group members that all contribute in chick raising (Rowley and Russell 1997). Red-backed fairywrens are no exception, as they routinely (though not always) have auxiliary helpers that help feed nestlings (Mulder et al. 1994). It may be that such assistance liberates females, at least in-part, from the need to compete over males that would provide the best parental care. Alternatively, as ornamented males appear to invest more in extra-pair copulations than in duller males (Webster et al. 2008; Dowling and Webster 2017), it may also be likely that ornamentation plays little-to-no role in choice à la parental care for red-backed fairywrens. In stark contrast, there is limited documentation (or anecdotal observation; JAJ pers. obs.) of helpers at the nest in the *M. a. moretoni* subspecies of white-shouldered fairywrens (Enbody et al. 2019), and helping behavior appears overall much rarer, relative to their Australian sister species. Interestingly, the unornamented subspecies of white-shouldered fairywren, *M. a. lorentzi*, is highly social (Boersma et al. 2022), live in large groups (Enbody et al. 2019), and anecdotally behave in a similar manner to red-backed fairywrens (i.e., similar social grouping and helper behavior). Futhermore, the unornamented female phenotype of *M. a. lorentzi* closely resembles that of female red-backed fairywrens and is thought to be the ancestral phenotype for the other, more ornamented white-shouldered fairywren subspecies (Enbody et al. 2022). A behavioral comparison between red-backed fairywrens and *M. a. lorentzi* would likely be a fruitful avenue of research, as these two species are share several life-history and mating system characteristics, despite the latitudinal distance between species.

In this preliminary study, we indirectly assessed the degree to which females within and between species compete over access to males of differing phenotypic quality as well as tested if commonly assumed behavioral differences between tropical and temperate birds were likely to be found in two sister species of fairywren. We failed to find support for our predictions that the most aggressive females would be mated to males of higher phenotypic quality, as measured by body size and plumage coloration, suggesting that female-female competition in these two species is likely over ecological resources when it occurs. We found that female white-shouldered fairywrens are on average more aggressive towards simulated same-sex rivals than females in their sister species, red-backed fairywrens, consistent with the idea that tropical birds experience an elevated competitive environment. This study raises as many questions as it answers about associations between female aggression and male phenotype (or quality), but it represents an important step in addressing these key questions. Together with the demonstration of the utility of mirror trials for studies of this sort that we have provided, we hope this work will spur further research into this current knowledge gap.

## Acknowledgments

In Papua New Guinea, our research would not have been possible without the logistical and in-field support from D. Nason and S. Ketaloya as well as the hospitality offered to us by the residents of Podagha Village (Milne Bay Province). We thank the Milne Bay provincial government for permits and permissions, and the National Research Institute for their assistance in acquiring country-level research permits (#99902100765) and visas. For Australia, we thank M. Webster for logistical support and Southeast Queensland Water for access to our field site. We respectfully acknowledge the Aboriginal peoples and Torres Strait Islander peoples as the Traditional Owners and Custodians of the land on which our research took place and pay our respects to the Elders both past and present.

Our study was funded by the National Science Foundation (IOS-1354133) awarded to JK, an Australian Department of Education and Training Endeavour Research Leadership fellowship (JAJ), the American Ornithological Society (JAJ), the American Philosophical Society (JAJ), the Department of Ecology and Evolutionary Biology of Tulane University (JAJ), Birds Queensland (WEF) and a Hermon Slade Project Grant (HS15/1) (WEF). Funders have had no influence on the content of the current manuscript nor do they require approval of the final manuscript to be published.

## References

Ah-King M. 2022. A female turn in bird research. In: The female turn: How evolutionary science shifted perceptions about females. Singapore: Palgrave Macmillan. p. 127–167.

Baldassarre DT, Webster M. 2013. Experimental evidence that extra-pair mating drives asymmetrical introgression of a sexual trait. Proc R Soc. 280:1–7. doi:10.1098/rspb.2013.2175. http://rspb.royalsocietypublishing.org/content/280/1771/20132175.short.

Beiko J, Lander R, Hampson E, Boon F, Cain DP. 2004. Contribution of sex differences in the acute stress response to sex differences in water maze performance in the rat. Behav Brain Res. 151(1–2):239–253. doi:10.1016/j.bbr.2003.08.019.

Boersma J, Jones JA, Enbody ED, Welklin JF, Ketaloya S, Nason D, Karubian J, Schwabl H. 2022. Hormones and behavior male white-shouldered fairywrens (Malurus alboscapulatus) elevate androgens greater when courting females than during territorial challenges. Horm Behav. 142:105158. doi:10.1016/j.yhbeh.2022.105158. 10.1016/j.yhbeh.2022.105158.x

Bro-Jørgensen J. 2002. Overt female mate competition and preference for central males in a lekking antelope. Proc Natl Acad Sci. 99(14):9290–9293. doi:10.1073/pnas.142125899.

Brouwer L, van de Pol M, Aranzamendi NH, Bain G, Baldassarre DT, Brooker LC, Brooker MG, Colombelli-Négrel D, Enbody E, Gielow K, et al. 2017. Multiple hypotheses explain variation in extra-pair paternity at different levels in a single bird family. Mol Ecol. 26(23):6717–6729. doi:10.1111/mec.14385.

Cain KE. 2014. Mates of competitive females: The relationships between female aggression, mate quality, and parental care. Adv Zool. 2014:319567. doi:10.1155/2014/319567.

Cain KE, Ketterson ED. 2012. Competitive females are successful females; phenotype, mechanism and selection in a common songbird. Behav Ecol Sociobiol. 66(2):241–252. doi:10.1007/s00265-011-1272-5. [accessed 2013 Oct 1]. http://www.pubmedcentral.nih.gov/articlerender.fcgi?artid=3278083&tool=pmcentrez&rendertype=abstract.

Cain KE, Ketterson ED. 2013. Costs and benefits of competitive traits in females: aggression, maternal care and reproductive success. PLoS One. 8(10). doi:10.1371/journal.pone.0077816.

Cain KE, Rosvall KA. 2014. Next steps for understanding the selective relevance of female-female competition. Front Ecol Evol. 2:32. doi:10.3389/fevo.2014.00032.

Catchpole CK, Slater PJB. 2008. Bird song: Biological themes and variations. Second Edi. Cambridge, UK: Cambridge University Press. https://www.cambridge.org/9780521872423.

Clutton-Brock TH. 2007. Sexual selection in males and females. Science (80-). 318:1882–1885. doi:10.1126/science.1133311.

Cockrem JF, Silverin B. 2002. Variation within and between birds in corticosterone responses of great tits (Parus major). Gen Comp Endocrinol. 125:197–206. doi:10.1006/gcen.2001.7750.

Crook JH. 1972. Sexual selection, dimorphism, and social organization in the primates. In: Campbell BG, editor. Sexual Selection and the Descent of Man. Chicago, IL: Aldine. p. 213– 281.

Cuthill IC. 2006. Color perception. In: Hill GE, McGraw. KJ, editors. Bird Coloration I: Mechanisms and Measurements. Cambridge, MA: Harvard University Press. p. 3–40.

Dale J, Dey C, Delhey K, Kempenaers B, Valcu M. 2015. The effects of life-history and social selection on male and female plumage coloration. Nature. 527(7578):367–370. doi:10.1038/nature15509. 10.1038/nature15509.

Delhey K, Delhey V, Kempenaers B, Peters A. 2015. A practical framework to analyze variation in animal colors using visual models. Behav Ecol. 26(2):367–375. doi:10.1093/beheco/aru198.

Delhey K, Hall M, Kingma SA, Peters A. 2013. Increased conspicuousness can explain the match between visual sensitivities and blue plumage colours in fairy-wrens. Proc R Soc B. 280(1750):20121771. doi:10.1098/rspb.2012.1771.

Dickens MJ, Earle KA, Romero LM. 2009. Initial transference of wild birds to captivity alters stress physiology. Gen Comp Endocrinol. 160(1):76–83. doi:10.1016/j.ygcen.2008.10.023. 10.1016/j.ygcen.2008.10.023.

Dowling J, Webster MS. 2017. Working with what you’ve got: unattractive males show greater mate-guarding effort in a duetting songbird. Biol Lett. 13(1):20160682. doi:10.1098/rsbl.2016.0682.

Enbody ED, Boersma J, Jones JA, Chatfield MWH, Ketaloya S, Nason D, Baldassarre DT, Hazlehurst J, Gowen O, Schwabl H, et al. 2019. Social organisation and breeding biology of the white-shouldered fairywren (Malurus alboscapulatus). Emu. 119(3):274–285. doi:10.1080/01584197.2019.1595663. 10.1080/01584197.2019.1595663.

Enbody ED, Sin SYW, Boersma J, Edwards S V., Ketaloya S, Schwabl H, Webster MS, Karubian J. 2022. The evolutionary history and mechanistic basis of female ornamentation in a tropical songbird. Evolution (N Y). 76(8):1720–1736. doi:10.1111/evo.14545.

Endler JA, Mielke PW. 2005. Comparing entire colour patterns as birds see them. Biol J Linn Soc. 86(4):405–431. doi:10.1111/j.1095-8312.2005.00540.x. http://doi.wiley.com/10.1111/j.1095-8312.2005.00540.x.

Fedy BC, Stutchbury BJM. 2005. Territory defence in tropical birds: Are females as aggressive as males? Behav Ecol Sociobiol. 58(4):414–422. doi:10.1007/s00265-005-0928-4.

Fischer S, Duffield C, Davidson AJ, Bolton R, Hurst JL, Stockley P. 2023. Fitness costs of female competition linked to resource defense and relatedness of competitors. Am Nat. 201:256– 268. doi:10.1086/722513.

Gallup Jr GG. 1968. Mirror-image stimulation. Psychol Bull. 70(6):782–793. doi:10.1037/h0026777. http://psycnet.apa.org/psycinfo/1969-06577-001.

Griffith SC. 2019. Cooperation and coordination in socially monogamous birds: Moving away from a focus on sexual conflict. Front Ecol Evol. 7:1–15. doi:10.3389/fevo.2019.00455.

Hart NS. 2001. The visual ecology of avian photoreceptors. Prog Retin Eye Res. 20:675–703. doi:10.1016/S1350-9462(01)00009-X.

Hasselquist D, Bensch S, von Schantz T. 1996. Correlation between male song repertoire, extra-pair paternity and offspring survival in the great reed warbler. Nature. 3181(6579):229–232. doi:10.1038/381229a0. https://www.nature.com.

Heinsohn R, Legge S, Endler JA. 2005. Evolution: Extreme reversed sexual dichromatism in a bird without sex role reversal. Science (80-). 309(5734):617–619. doi:10.1126/science.1112774. http://www.sciencemag.org/cgi/content/abstract/309/5734/617.

Hurley LL, Rowe M, Griffith SC. 2020. Reproductive coordination breeds success: The importance of the partnership in avian sperm biology. Behav Ecol Sociobiol. 74:3. doi:10.1007/s00265-019-2782-9.

Ihle M, Kempenaers B, Forstmeier W. 2015. Fitness benefits of mate choice for compatibility in a socially monogamous species. PLoS Biol. 13(9):1–21. doi:10.1371/journal.pbio.1002248.

Jawor JM, Young R, Ketterson ED. 2006. Females competing to reproduce: Dominance matters but testosterone may not. Horm Behav. 49(3):362–368. doi:10.1016/j.yhbeh.2005.08.009.

Jones JA, Boersma J, Karubian J. 2022. Female aggression towards same-sex rivals depends on context in a tropical songbird. Behav Processes. 202:104735. doi:10.1016/j.beproc.2022.104735. 10.1016/j.beproc.2022.104735.

Jones JA, Boersma J, Liu J, Nason D, Ketaloya S, Karubian J. 2022. Female ornamentation does not predict aggression in a tropical songbird. Behav Ecol Sociobiol. 76:57. doi:10.1007/s00265-022-03165-x.

Kappeler PM, Benhaiem S, Fichtel C, Fromhage L, Höner OP, Jennions MD, Kaiser S, Krüger O, Schneider JM, Tuni C, et al. 2022. Sex roles and sex ratios in animals. Biol Rev. 6. doi:10.1111/brv.12915.

Karubian J. 2002. Costs and benefits of variable breeding plumage in the red-backed fairy-wren. Evolution (N Y). 56(8):1673–1682. doi:10.1554/0014-3820(2002)056[1673:CABOVB]2.0.CO;2.

Karubian J. 2008. Changes in breeding status are associated with rapid bill darkening in male red-backed fairy-wrens Malurus melanocephalus. J Avian Biol. 39(1):81–86. doi:10.1111/j.0908-8857.2008.04161.x.

Karubian J. 2013. Female ornamentation in Malurus fairy-wrens: A hidden evolutionary gem for understanding female perspectives on social and sexual selection. Emu. 113(3):248–258. doi:10.1071/MU12093.

Karubian J, Lindsay WR, Schwabl H, Webster MS. 2011. Bill coloration, a flexible signal in a tropical passerine bird, is regulated by social environment and androgens. Anim Behav. 81(4):795–800. doi:10.1016/j.anbehav.2011.01.012. 10.1016/j.anbehav.2011.01.012.

Karubian J, Sillett TS, Webster MS. 2008. The effects of delayed plumage maturation on aggression and survival in male red-backed fairy-wrens. Behav Ecol. 19(3):508–516. doi:10.1093/beheco/arm159.

Karubian J, Swaddle JP, Varian-Ramos CW, Webster MS. 2009. The relative importance of male tail length and nuptial plumage on social dominance and mate choice in the red-backed fairy-wren Malurus melanocephalus: Evidence for the multiple receiver hypothesis. J Avian Biol. 40(5):559–568. doi:10.1111/j.1600-048X.2009.04572.x.

Kraaijeveld K, Kraaijeveld-Smit FJL, Komdeur J. 2007. The evolution of mutual ornamentation. Anim Behav. 74(4):657–677. doi:10.1016/j.anbehav.2006.12.027.

Kraft FL, Forštová T, Utku Urhan A, Exnerová A, Brodin A. 2017. No evidence for self-recognition in a small passerine, the great tit (Parus major) judged from the mark/mirror test. Anim Cogn. 20(6):1049–1057. doi:10.1007/s10071-017-1121-7.

Kunkel P. 1974. Mating systems of tropical birds: The effects of weakness or absence of external reproduction□timing factors, with special reference to prolonged pair bonds. Z Tierpsychol. 34(3):265–307. doi:10.1111/j.1439-0310.1974.tb01802.x.

Langston NE, Freeman S, Rohwer SA, Gori D. 1990. The evolution of female body size in red-winged blackbirds: The effects of timing of breeding, social competitiong, and reproductive energetics. Evolution (N Y). 44(7):1764–1779. doi:10.1111/j.1558-5646.1990.tb05247.x.

Leitão AV., Hall ML, Delhey K, Mulder RA. 2019. Female and male plumage colour signals aggression in a dichromatic tropical songbird. Anim Behav. 150:285–301. doi:10.1016/j.anbehav.2019.01.025. https://linkinghub.elsevier.com/retrieve/pii/S0003347219300375.

Lipshutz SE, Rosvall KA. 2021. Nesting strategy shapes territorial aggression but not testosterone: A comparative approach in female and male birds. Horm Behav. 133:104995. doi:10.1016/j.yhbeh.2021.104995. 10.1016/j.yhbeh.2021.104995.

Lyon BE, Montgomerie R. 2012. Sexual selection is a form of social selection. Philos Trans R Soc B. 367(1600):2266–2273. doi:10.1098/rstb.2012.0012.

Mays HL, Hill GE. 2004. Choosing mates: Good genes versus genes that are a good fit. Trends Ecol Evol. 19(10):554–559. doi:10.1016/j.tree.2004.07.018.

Mulder RA, Dunn PO, Cockburn A, Lazenby-Cohen KA, Howell MJ. 1994. Helpers liberate female fairy-wrens from constraints on extra-pair mate choice. Proc R Soc B. 255(1344):223– 229. doi:10.1098/rspb.1994.0032.

Murphy TG, Hernández-Muciño D, Osorio-Beristain M, Montgomerie R, Omland KE. 2009. Carotenoid-based status signaling by females in the tropical streak-backed oriole. Behav Ecol. 20(5):1000–1006. doi:10.1093/beheco/arp089.

Ödeen A, Håstad O. 2013. The phylogenetic distribution of ultraviolet sensitivity in birds. BMC Evol Biol. 13:36. doi:10.1186/1471-2148-13-36.

Ödeen A, Pruett-Jones S, Driskell AC, Armenta JK, Håstad O. 2012. Multiple shifts between violet and ultraviolet vision in a family of passerine birds with associated changes in plumage coloration. Proc R Soc B. 279(1732):1269–1276. doi:10.1098/rspb.2011.1777.

Oliveira RF, Carneiro LA, Canário A V. 2005. No hormonal response in tied fights. Nature. 437:207–208. doi:10.1038/437207a.

Oliveira RF, Simes JM, Teles MC, Oliveira CR, Becker JD, Lopes JS. 2016. Assessment of fight outcome is needed to activate socially driven transcriptional changes in the zebrafish brain. Proc Natl Acad Sci. 113(5):E654–E661. doi:10.1073/pnas.1514292113.

Olsson P, Lind O, Kelber A. 2018. Chromatic and achromatic vision: parameter choice and limitations for reliable model predictions. Behav Ecol. 29(2):273–282. doi:10.1093/beheco/arx133. http://fdslive.oup.com/www.oup.com/pdf/production_in_progress.pdf.

Pagani-Núñez E, Senar JC. 2014. Are colorful males of great tits Parus major better parents? Parental investment is a matter of quality. Acta Oecologica. 55:23–28. doi:10.1016/j.actao.2013.11.001. 10.1016/j.actao.2013.11.001.

Préault M, Chastel O, Cézilly F, Faivre B. 2005. Male bill colour and age are associated with parental abilities and breeding performance in blackbirds. Behav Ecol Sociobiol. 58(5):497–505. doi:10.1007/s00265-005-0937-3.

Prior H, Schwarz A, Güntürkün O. 2008. Mirror-induced behavior in the magpie (Pica pica): Evidence of self-recognition. Plos Biol. 6(8):e202. doi:10.1371/journal.Citation.

Pryke SR. 2007. Fiery red heads: Female dominance among head color morphs in the Gouldian finch. Behav Ecol. 18(3):621–627. doi:10.1093/beheco/arm020.

Reudink MW, Studds CE, Marra PP, Kurt Kyser T, Ratcliffe LM. 2009. Plumage brightness predicts non-breeding season territory quality in a long-distance migratory songbird, the American redstart Setophaga ruticilla. J Avian Biol. 40(1):34–41. doi:10.1111/j.1600-048X.2008.04377.x.

Rosvall KA. 2008. Sexual selection on aggressiveness in females: Evidence from an experimental test with tree swallows. Anim Behav. 75(5):1603–1610. doi:10.1016/j.anbehav.2007.09.038.

Rosvall KA. 2011. Intrasexual competition in females: Evidence for sexual selection? Behav Ecol. 22(6):1131–1140. doi:10.1093/beheco/arr106.

Rowe M, McGraw KJ. 2008. Carotenoids in the seminal fluid of wild birds: Interspecific variation in fairy-wrens. Condor. 110(4):694–700. doi:10.1525/cond.2008.8604.

Rowe M, Pruett-Jones S. 2011. Sperm competition selects for sperm quantity and quality in the Australian Maluridae. PLoS One. 6(1):e15720. doi:10.1371/journal.pone.0015720.

Rowe M, Pruett-Jones S. 2013. Extra-pair paternity, sperm competition and their evolutionary consequences in the Maluridae. Emu. 113(3):218–231. doi:10.1071/MU12084.

Rowley I, Russell E. 1997. Bird families of the world: Fairy-wrens and Grasswrens. Oxford, United Kingdom: Oxford University Press.

Slagsvold T, Lifjeld JT. 1996. Polygyny in birds: The role of competition between females for male parental care. Am Nat. 143(1):59–94. doi:10.1086/285596.

Slater PJB, Mann NI. 2004. Why do the females of many bird species sing in the tropics? J Avian Biol. 35:289–294. doi:10.1111/j.0908-8857.2004.03392.x.

Stockley P, Bro-Jørgensen J. 2011. Female competition and its evolutionary consequences in mammals. Biol Rev. 86(2):341–366. doi:10.1111/j.1469-185X.2010.00149.x.

Stockley P, Campbell A. 2013. Female competition and aggression: Interdisciplinary perspectives. Philos Trans R Soc B. 368:20130073. doi:10.1098/rstb.2013.0073.

Stutchbury BJM, Morton ES. 2023. Behavioral ecology of tropical birds. 2nd editio. Academic Press.

Swaddle JP, Karubian J, Pruett-Jones S. 2000. A novel evolutionary pattern of reversed sexual dimorphism in fairy wrens: Implications for sexual selection. Behav Ecol. 11(3):345–349. doi:10.1093/beheco/11.3.345.

Team RC. 2021. R: A language and environment for statistical computing. R Foundation for Statistical Computing, Vienna, Austria. https://www.R-project.org/.

Teles MC, Dahlbom SJ, Winberg S, Oliveira RF. 2013. Social modulation of brain monoamine levels in zebrafish. Behav Brain Res. 253:17–24. doi:10.1016/j.bbr.2013.07.012. 10.1016/j.bbr.2013.07.012.

Tobias JA, Montgomerie R, Lyon BE. 2012. The evolution of female ornaments and weaponry: Social selection, sexual selection and ecological competition. Philos Trans R Soc B. 367(1600):2274–2293. doi:10.1098/rstb.2011.0280.

Trivers RL. 1972. Parental investment and sexual selection. In: Sexual Selection and the Descent of Man. Chicago, IL: Aldine Publishing Compnay. p. 136–179.

Tuttle EM, Pruett-Jones S, Webster MS. 1996. Cloacal protuberances and extreme sperm production in Australian fairy-wrens. Proc R Soc B Biol Sci. 263(1375):1359–1364. doi:10.1098/rspb.1996.0199.

Vorobyev M, Osorio D. 1998. Receptor noise as a determinant of colour thresholds. Proc R Soc B. 265:351–358. doi:10.1098/rspb.1998.0302.

Vorobyev M, Osorio D, Bennett AT, Marshall NJ, Cuthill IC. 1998. Tetrachromacy, oil droplets and bird plumage colours. J Comp Physiol A. 183:621–633. doi:10.1007/s003590050286.

Wang D, Forstmeier W, Kempenaers B. 2017. No mutual mate choice for quality in zebra finches: Time to question a widely held assumption. Evolution (N Y). 71(11):2661–2676. doi:10.1111/evo.13341.

Webster MS, Varian CW, Karubian J. 2008. Plumage color and reproduction in the red-backed fairy-wren: Why be a dull breeder? Behav Ecol. 19(3):517–524. doi:10.1093/beheco/arn015.

West-Eberhard MJ. 1979. Sexual selection, social competition, and evolution. Proc Am Philos Soc. 123(4):222–234.

Westneat DF, Sargent RC. 1996. Sex and parenting: The effects of sexual conflict and parentage on parental strategies. Trends Ecol Evol. 11(2):87–91. doi:10.1016/0169-5347(96)81049-4. http://www.sciencedirect.com/science/article/B6VJ1-3WJG1XR-6G/2/521cf5f6aa7339906ef1800b9581b44d.

Whiteman EA, Côté IM. 2003. Social monogamy in the cleaning goby Elacatinus evelynae: Ecological constraints or net benefit? Anim Behav. 66(2):281–291. doi:10.1006/anbe.2003.2200.

Yasukawa K, Searcy WA. 1982. Aggression in female red-winged blackbirds: A strategy to ensure male parental investment. Behav Ecol Sociobiol. 11(1):13–17. doi:10.1007/BF00297660.

